# Evaluating the contribution of cell-type specific alternative splicing to variation in lipid levels

**DOI:** 10.1101/659326

**Authors:** K.A.B. Gawronski, W. Bone, Y. Park, E. Pashos, X. Wang, W. Yang, D. Rader, K. Musunuru, B. Voight, C. Brown

## Abstract

**Background:** Genome-wide association studies have identified 150+ loci associated with lipid levels. However, the genetic mechanisms underlying most of these loci are not well-understood. Recent work indicates that changes in the abundance of alternatively spliced transcripts contributes to complex trait variation. Consequently, identifying genetic loci that associate with alternative splicing in disease-relevant cell types and determining the degree to which these loci are informative for lipid biology is of broad interest.

**Methods and Results:** We analyze gene splicing in 83 sample-matched induced pluripotent stem cell (iPSC) and hepatocyte-like cell (HLC) lines (n=166), as well as in an independent collection of primary liver tissues (n=96). We observe that transcript splicing is highly cell-type specific, and the genes that are differentially spliced between iPSCs and HLCs are enriched for metabolism pathway annotations. We identify 1,381 HLC splicing quantitative trait loci (sQTLs) and 1,462 iPSC sQTLs and find that sQTLs are often shared across cell types. To evaluate the contribution of sQTLs to variation in lipid levels, we conduct colocalization analysis using lipid genome-wide association data. We identify 19 lipid-associated loci that colocalize either with an HLC expression quantitative trait locus (eQTL) or sQTL. Only one locus colocalizes with both an sQTL and eQTL, indicating that sQTLs contribute information about GWAS loci that cannot be obtained by analysis of steady-state gene expression alone.

**Conclusions:** These results provide an important foundation for future efforts that use iPSC and iPSC-derived cells to evaluate genetic mechanisms influencing both cardiovascular disease risk and complex traits in general.

## Introduction

Genome-wide association studies have identified hundreds of loci associated with plasma lipid levels, an important set of predictive and causal risk factors for cardiovascular disease.^1^ The majority of variants associated with these traits are found in the non-coding genome and are hypothesized to mechanistically influence complex traits through changes in gene expression. For example, at *SORT1*, an established locus associated with both LDL cholesterol levels and coronary heart disease, a functional noncoding variant creates a novel C/EBP binding site, leading to changes in hepatic *SORT1* expression and in turn, changes in circulating LDL cholesterol levels.^2^ Motivated by this and other examples, significant effort has been dedicated to identifying variants associated with changes in gene expression, or expression quantitative trait loci (eQTLs) that are also associated with changes in plasma lipid levels.

Recent research indicates that variants associated with changes in the proportion of alternatively spliced transcript isoforms (*i.e.*, sQTLs) can provide another contributing mechanism underlying complex traits, and may be as informative as total gene expression quantitative trait loci in some cases.^3,4^ For example, sQTLs discovered in lymphoblast cell lines are enriched in multiple sclerosis GWAS disease loci, and other studies report similar findings for sQTL enrichment in GWAS for schizophrenia and type-2 diabetes.^4–6^ Variants affecting splicing have also been linked to systemic lupus erythematosus and fatty acid metabolism.^7,8,9^ Splicing is an attractive mechanism for study because it can be targeted with antisense oligonucleotides, several of which are currently undergoing clinical trials to treat diseases resulting from aberrant splicing such as Duchenne’s Muscular Dystrophy and Spinal Muscular Atrophy.^10^ Finally, splicing provides an even finer level of detail on biological mechanism that cannot always be inferred through bulk expression analysis alone. For example, the Alzheimer’s disease-associated gene *ABCA7* has both an eQTL and sQTL association. The sQTL is hypothesized to be the causal signal, as it is associated with the production of a non-functional transcript.^11,12^

Functional studies of eQTLs have been hindered by the time and energy required to identify and then mechanistically characterize the QTLs in model systems. QTL discovery efforts in primary tissues have been highly productive, but have several important drawbacks, such as the reliance on heterogeneous post-mortem tissue collection and the difficulty of interrogating phenotypes in tissues. However, it is now possible to generate individual-specific, renewable, induced pluripotent stem cell (iPSC) lines, which can be used to both identify and characterize QTLs in specific cell types. iPSCs can be differentiated into a variety of cell types, including hepatocyte-like cells (HLCs).^13^ Given the liver’s importance in the synthesis and uptake of lipids,^14^ using HLCs as a model to understand the mechanisms underlying genetic associations with lipid levels is of particular interest. We and others have previously demonstrated the utility of these cell models by identifying and characterizing eQTLs in these HLCs.^15,16^

Therefore, this report has two goals, namely: (i) to evaluate the feasibility of using HLC/iPSC lines for sQTL discovery, and (ii) to determine whether the sQTLs discovered are informative for our understanding of lipid biology. Therefore, we mapped sQTLs in 83 individual-matched iPSC lines and iPSC-derived HLCs, on which we conducted genotyping and paired-end RNA-sequencing. To evaluate the degree to which the genetic control of splicing information compares to total gene expression, we also perform a variety of analyses evaluating differential splicing, differential expression, and colocalization analysis with both HLC sQTLs and HLC eQTLs. We then highlight two loci where the colocalization of GWAS and sQTL results provide detail as to the causal mechanism underlying the variant-trait association.

## Results

### iPSC and HLC samples have distinct profiles of gene expression and alternative splicing

We first sought to compare the patterns of alternative splicing and gene expression between iPSCs and HLCs. After normalization and standardization procedures (**Methods**), we applied Leafcutter to identify genes with differences in proportions of alternatively spliced transcripts. 9,484 genes out of 12,213 genes tested are differentially spliced between iPSC and HLC samples (false discovery rate, FDR <5%). The first two principal components of the quantile normalized and standardized splicing proportions clearly separate the samples by cell type (Figure 1a). Gene ontology analysis also demonstrated that differentially spliced genes that had at least a 10% splicing difference across cell types were enriched for metabolism-relevant KEGG pathways such as insulin secretion (Figure 1b).

**Figure 1.**
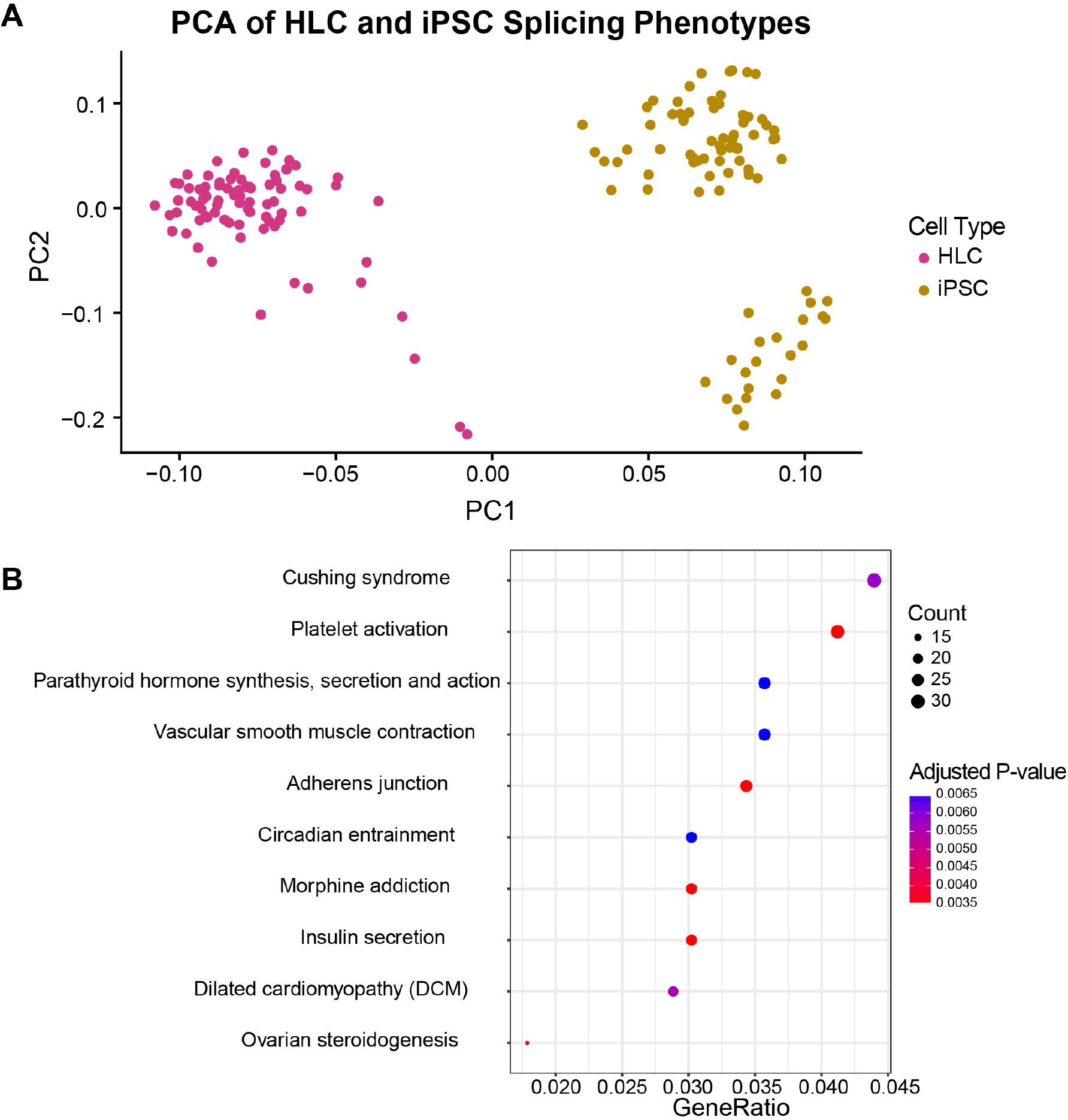
Differential splicing analysis: a) PCA of normalized splicing phenotypes; b) enrichment of KEGG pathways for differentially spliced genes with a delta PSI ≥ 0.1. P-values were adjusted for multiple testing using the Benjamini and Hochberg correction.

We next compared patterns of gene expression between the iPSC and HLC lines (**Methods**). Virtually all (22,312 out of 23,857) tested genes were differentially expressed between iPSCs and HLCs (FDR < 5%), with 6,045 having an absolute log_2_ fold change of 2 or more. Differentially expressed genes that were more highly expressed in HLCs (relative to iPSCs) were enriched for KEGG pathways pertaining to liver function such as cholesterol metabolism, drug metabolism, and fat digestion and absorption (Supplementary Figure 1c). The first two principal components obtained from the normalized expression profiles for the iPSCs and HLCs clearly separated samples by cell type (Supplementary Figure 1a). In sum, we found substantial differences in alternative splicing and gene expression between iPSC and HLCs, and the top differentially expressed or alternatively spliced genes were enriched for liver, lipid, and metabolism relevant pathways. These results indicate that the successful differentiation of the iPSCs into HLCs results in broad changes in steady state transcriptional and post-transcriptional regulation.

### Thousands of sQTLs identified across iPSC and HLC samples

To identify genetic variants associated with differences in the proportion of alternatively spliced transcripts (i.e., sQTLs), we conducted sQTL discovery scans in both cell types using QTLtools (**Methods**). We identify 1,381 sQTLs in HLCs and 1,462 sQTLs in iPSCs (FDR < 5%, **Supplementary Table 1**). iPSC sQTLs correspond to 1,444 unique SNPs and 1,462 unique genes, while HLC sQTLs correspond to 1,365 unique SNPs and 1,381 unique genes.

To evaluate the extent to which iPSC and HLC sQTLs are discoverable in primary liver tissue, we performed replication analysis in GTEx v6 primary liver samples (n=96).^17^ As one would expect, we observed more biologically similar cell types had more similar sQTL profiles. In comparison to iPSC sQTLs, HLC sQTLs were more likely to be replicated at the gene-level in primary liver tissue (π_1_ = 0.82 for HLC sQTLs vs. π_1_ = 0.76 for iPSC sQTLs). These results demonstrate that sQTL discovery in iPSC and HLC samples is feasible, and that both iPSC and HLC sQTLs replicate in primary liver tissue, with HLC sQTLs replicating at a higher degree than iPSCs.

### sQTLs are found near their associated splice event and are enriched for splicing relevant annotations

Because alternative splicing involves protein regulatory machinery interacting with genomic elements near intron/exon boundaries, we expect true sQTL variants to be found predominantly in the regions near these boundaries. Thus, we assessed the genomic context of each sQTL variant and measured the distance between each variant and its associated splice event. The majority of lead sQTL variants (69%) are within 25kb of their associated splice event, despite the fact that the lead sentinel variant may not be the ultimate causal variant. We observe similar patterns for sQTLs that fall within the intronic region of their splice event (in between the two exons that delineate the excised intron). After binning to normalize for intron length, we observe that the windows that are closest to canonical splice sites (the 1st and 10th deciles from Figure 2) contain the highest numbers of sQTLs (Figure 2a, 2b). Finally, both sets of HLC and iPSC sQTLs are significantly enriched in the 11bp window encompassing ends of exons, extending into introns and in several other genomic annotations (Figure 2c). These observations demonstrate that our identified sQTLs are often found close enough to canonical splicing regulatory elements to facilitate fine-mapping of causal variants and genes.

**Figure 2.**
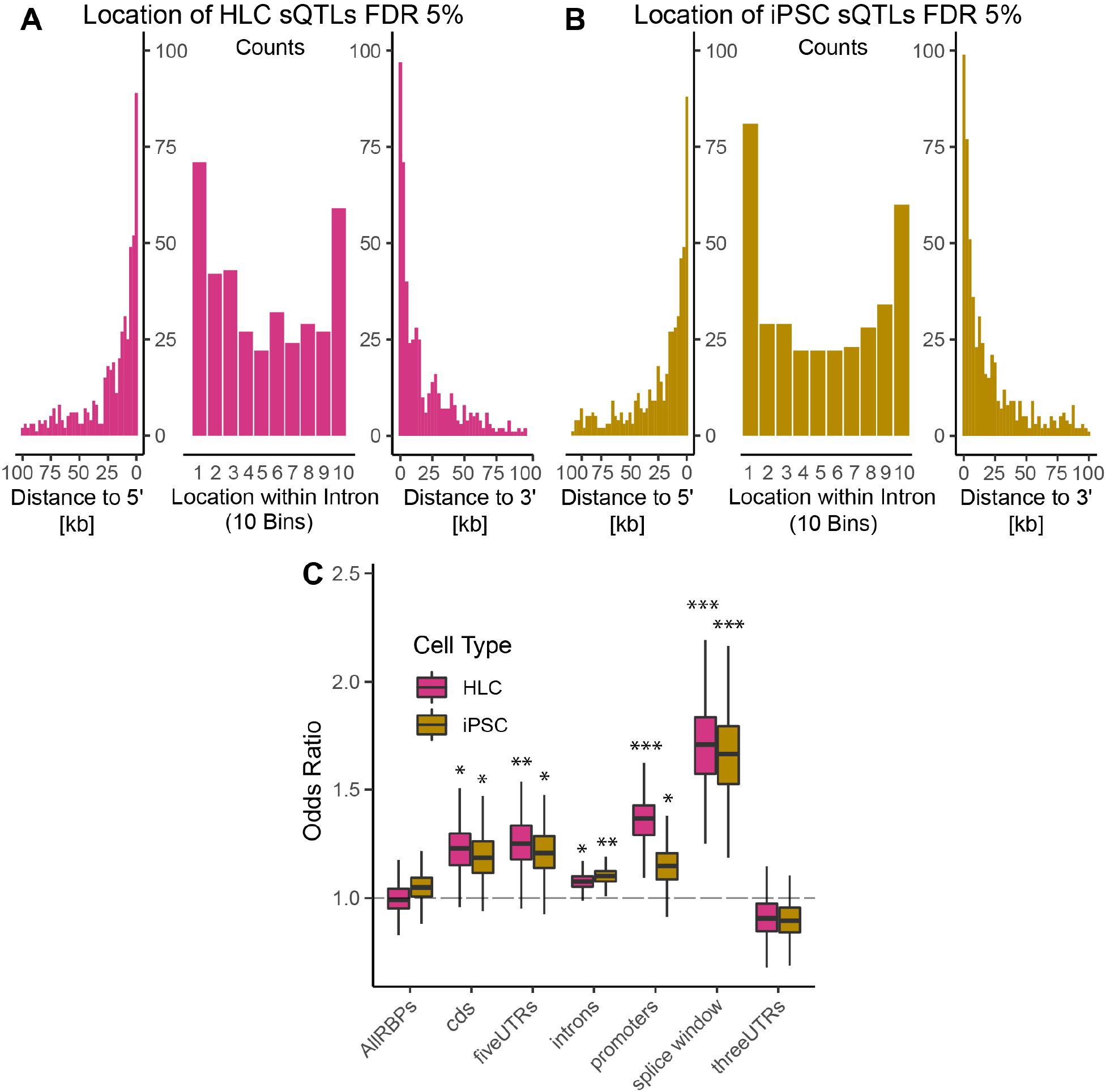
Characteristics of FDR <5% sQTLs: a) distance of iPSC sQTLs to splice event, b) distance of HLC sQTLs to splice event, c) enrichment of iPSC/HLC sQTLs in genomic annotations - boxplot whiskers calculated as interquartile range*1.5. ***P < 0.001; **P< 0.01; *P<0.05

We also conducted credible set fine-mapping analysis of the colocalized sQTLs to identify 95% credible set SNPs; 75% of the colocalized sQTLs have a credible set SNP located within the sQTL-associated gene. Median credible set sizes for iPSC and HLC sQTLs were comparable to that of iPSC and HLC eQTLs – median credible set size for sQTLs is 9 and 8 SNPs for iPSC and HLC samples respectively, while median credible set size for eQTLs is 11 and 9 SNPs for iPSC and HLC samples. This suggests that sQTLs will be no more difficult to fine-map and functionally characterize than eQTLs. These observations suggest that sQTLs contribute information pertaining to potential causal genes and molecular mechanisms that may not otherwise be captured by eQTLs.

### A majority of sQTLs are shared between iPSCs and HLCs

Previous research has indicated that sQTL effects are often shared across tissues.^18,19^ To evaluate the extent of sQTL sharing in our cell types, we examined the top one thousand sQTLs from the FDR < 5% sQTL set for both the iPSC and HLC samples, and used METASOFT to determine the extent to which sQTLs discovered in one cell type are found in the other. We find that if an alternatively spliced isoform is expressed in both cell types, the sQTL effect is observed in both cell types more than 90% of the time (Figure 3a, 3b). In contrast, 60-74% of eQTL effects are shared, appearing to be more cell-type restricted (Figure 3c, 3d). However, this difference may simply be due to statistical power for discovery. The power to detect sQTLs is lower than that of eQTLs, due to a higher burden of multiple testing correction (**Methods**). In GTEx, tissues with larger sample sizes report higher percentages of tissue-specific eQTLs compared to tissues with smaller sample sizes, indicating that power is correlated with tissue-specific eQTL discovery. The high percentage of shared sQTLs identified in this study may indicative of similar power limitations.

**Figure 3.**
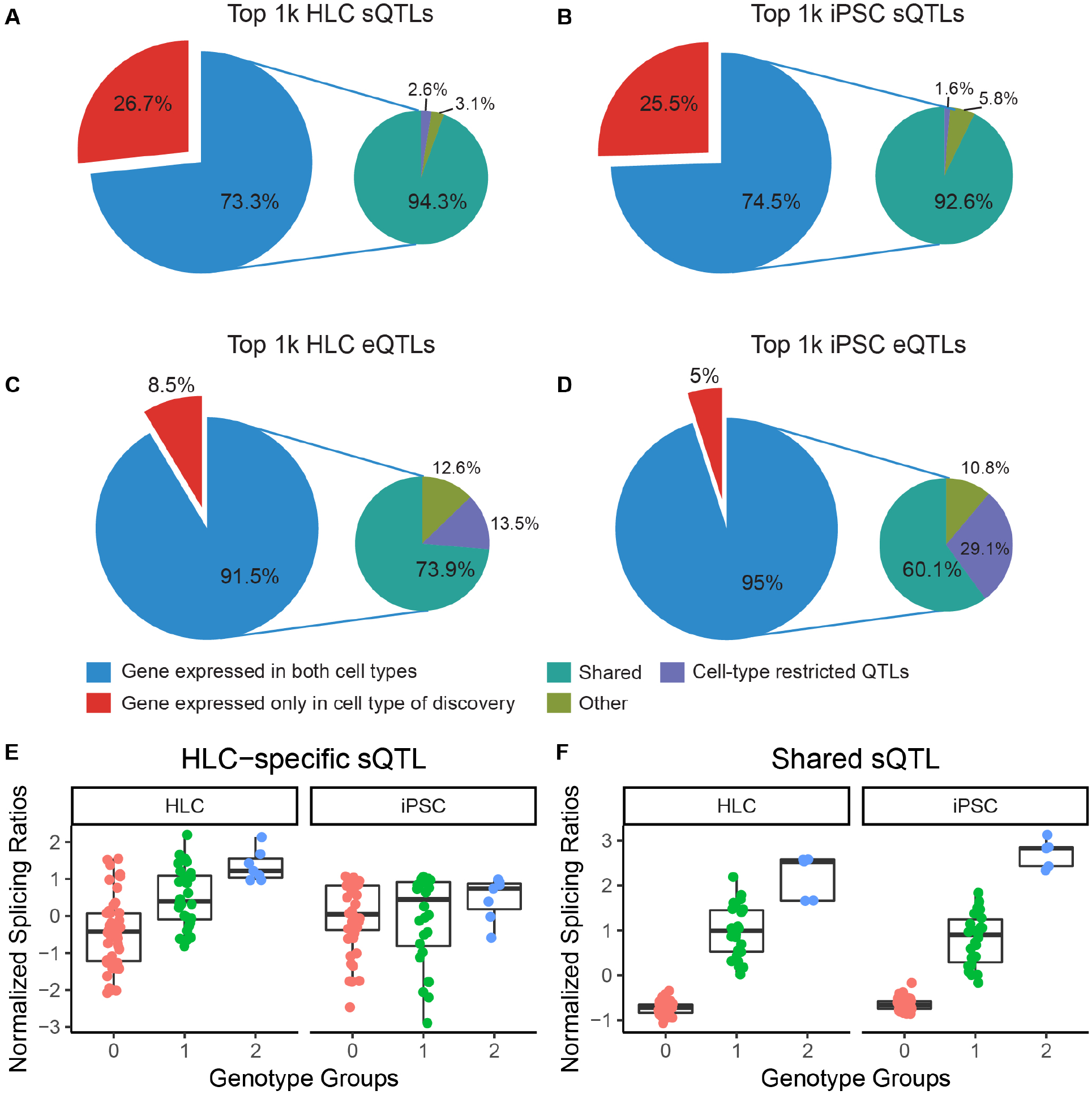
Percentage of the top one thousand: a) HLC sQTLs, b) iPSC sQTLs c) HLC eQTLs, d) iPSC eQTLs, e) sQTL that is present in HLCs only, f) sQTL that is identified in both HLCs and iPSCs.

### HLC sQTLs identify trait-relevant genes and mechanisms at GWAS lipid loci

To determine if our discovered sQTLs map to previously established associations for lipid traits in humans, we evaluated the probability that sQTL and GWAS signals share underlying causal architecture using a statistical process known as colocalization (as implemented by coloc R package).^20^ Colocalization analysis of HLC sQTLs (FDR < 5%) with genome-wide significant (p<5e-08) loci from the Global Lipids Genetics Consortium^1^ identified 5,3,4, and 5 HLC sQTLs colocalizing with HDL, LDL, TG, and TC loci respectively (using a cutoff of PP4/PP3+PP4 ≥0.9 and PP3+PP4 ≥0.8, Table 1).^21^ Colocalization analysis of FDR <5% HLC eQTLs identified 2, 5, 2, and 5 eQTL/GWAS colocalized loci for HDL, LDL, TG, and TC respectively (Table 2). Encouragingly, several of the colocalized sQTL genes have known roles in lipid biology (*APOC1*),^22^ and eQTLs previously shown to colocalize with GLGC traits and be functional for lipid levels are replicated (*ANGPTL3).*^15^ In general, the colocalized sQTLs and eQTLs are associated with different genes, a result that is in line with previous studies.^18^ Only a single gene colocalized for both an eQTL and sQTL, *RPAP2*.

**Table 1.**
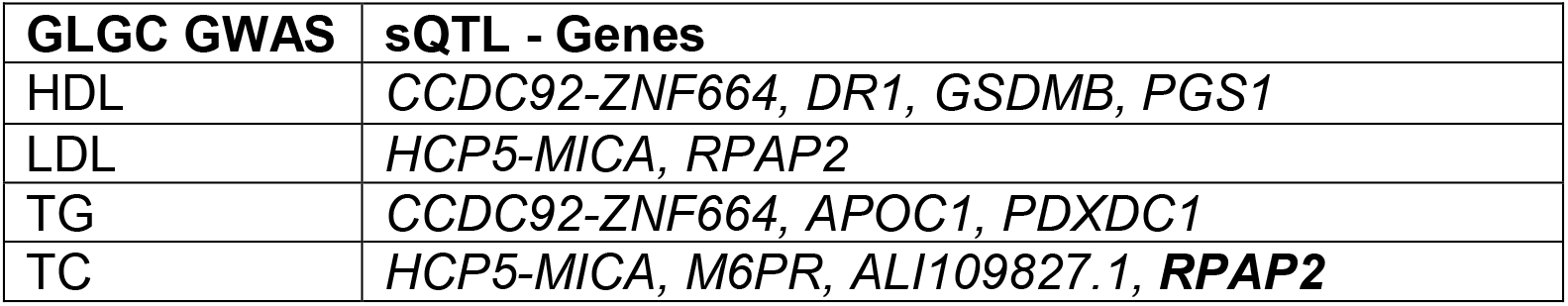
HLC sQTL Genes that Colocalize with GLGC Loci

| **GLGC GWAS** | **sQTL - Genes** |
| --- | --- |
| HDL | *CCDC92-ZNF664, DR1, GSDMB, PGS1* |
| LDL | *HCP5-MICA, RPAP2* |
| TG | *CCDC92-ZNF664, APOC1, PDXDC1* |
| TC | *HCP5-MICA, M6PR, ALI109827.1, **RPAP2*** |

**Table 2.** HLC eQTL Genes that Colocalize with GLGC Loci

| **GLGC GWAS** | **eQTL - Genes** |
| --- | --- |
| HDL | *RBM6, PGAP3* |
| LDL | *SPTY2D1, SYPL2, FRK, TBKBP1, **RPAP2*** |
| TG | *ANGPTL3, AL138847.1* |
| TC | *SPTY2D1, ANGPTL3, FRK, TBKBP1, AL138847.1* |

Of particular interest are the colocalized sQTLs mapping to *CCDC92* and *PGS1*. *PGS1* encodes an enzyme known as Phosphatidylglycerophosphate Synthase 1, which is involved in the synthesis of the anionic phospholipids phospatidylglycerol and cardiolipin. SNPs in *PGS1* are associated with changes in triglyceride levels during diabetes treatment.^23^ The second SNP in the credible set for this sQTL falls within the canonical splice site of the splicing event (as demonstrated by the red arrow in Figure 4d). Further, the differentially spliced intron for *PGS1* is associated with a transcript that is predicted to undergo nonsense-mediated decay (*PGS1-002*). The *CCDC92-ZNF664* locus has also been implicated in many different metabolic disorders, including type-2 diabetes and coronary heart disease.^24^ The lead sQTL SNP at this colocalized locus has been associated with variation in total cholesterol, metabolic syndrome, and waist circumference in previous studies.^24^

**Figure 4.**
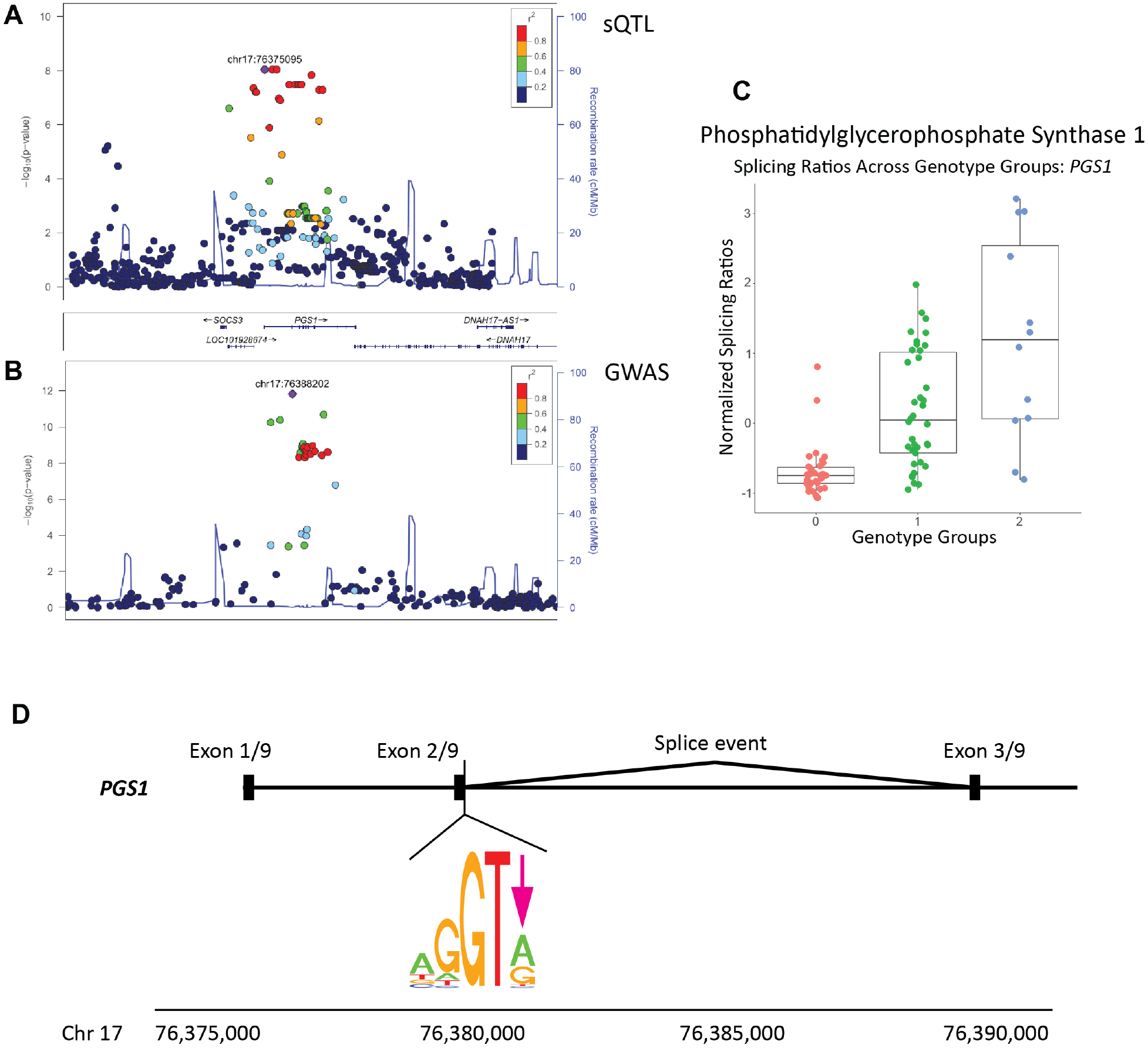
*PGS1* colocalization result: a) sQTL and b) GWAS locuszoom plots for *PGS1* locus, c) boxplots of normalized splicing ratios for sQTL across genotype groups, d) location of credible set SNP relative to associated exon-exon junction (indicated by red arrow). Position Weight Matrices used to create the sequence logo obtained from Abril et al. 2005.^47^

**Figure 5.**
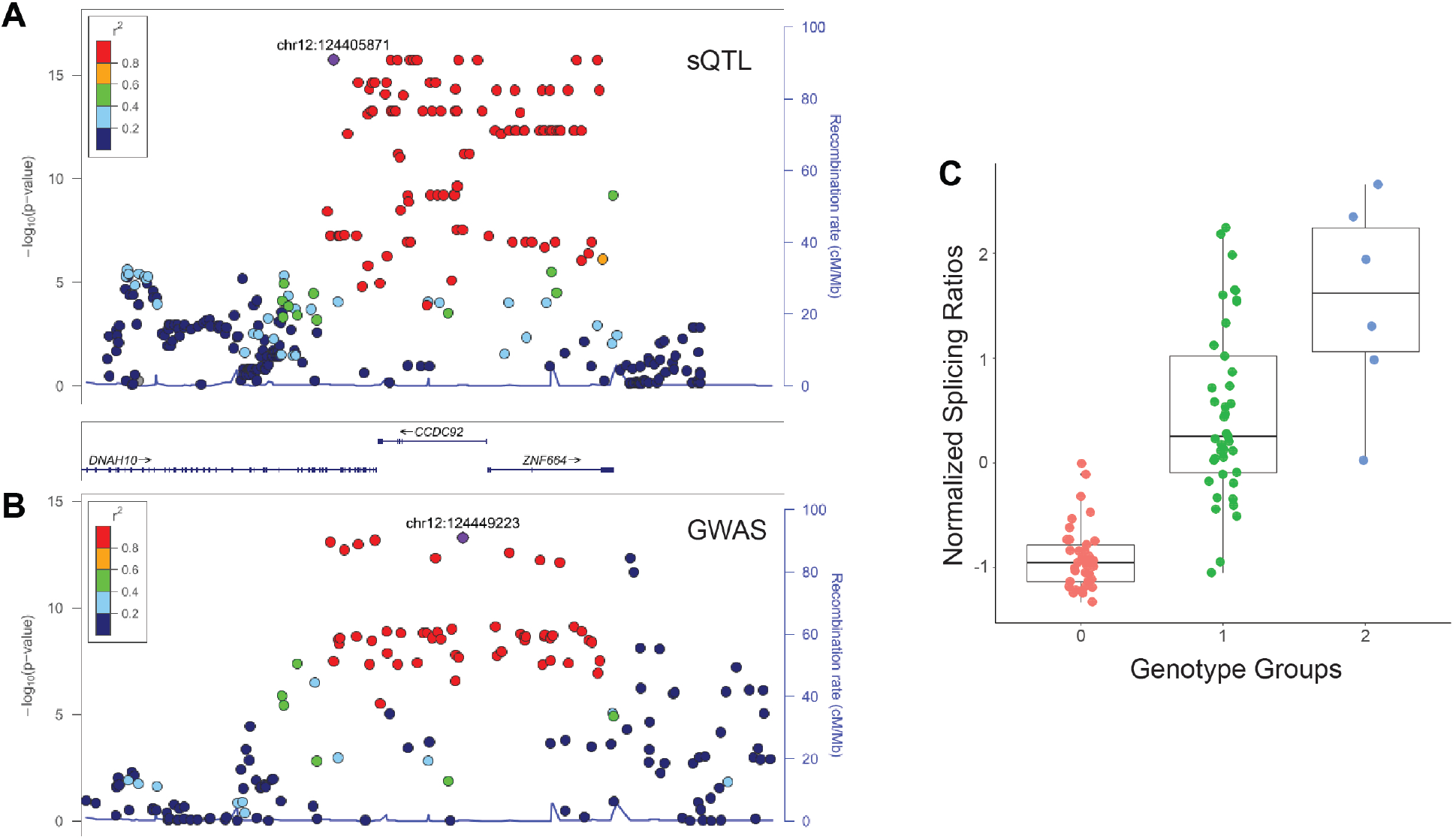
*CCDC92* colocalization result: a) sQTL and b) GWAS locuszoom plots for *CCDC92* locus, c) boxplots of normalized splicing ratios for sQTL across genotype groups for sQTL mapping to *CCDC92*.

## Discussion

In sum, we demonstrate the utility of using iPSC-derived hepatocyte-like cells for the identification of lipid-relevant sQTLs. Our sQTL scans identify thousands of sQTLs in both iPSC and HLC lines, and the lead sQTL variants are enriched near canonical splice sites and are located close to their associated splice event, indicating that fine-mapping the causal genes and variants for these sQTLs is feasible. Regarding the degree to which sQTLs are shared across cell-types, we find that if a spliced isoform is expressed across both cell types, the sQTL effect is also often shared. Although this finding is in line with previous sQTL analyses, it is may be due to the fact that lowly expressed splice events are often removed during more stringent filtering processes.

Our colocalization analysis of the HLC sQTLs with GWAS loci from the GLGC identifies several interesting genes and variants that underlie blood lipid level variation. Further, the fact that: 1) our HLC sQTLs generally colocalize with different GWAS loci than the HLC eQTLs, and 2) the colocalized sQTLs nominate different genes than the eQTLs, indicates that sQTLs likely provide information pertaining to the causal genes and molecular mechanisms underlying complex traits that may not otherwise be captured by eQTL analysis alone.

One limitation of this study is the fact that Leafcutter cannot distinguish between types of sQTL events and may miss variants associated with alternative transcription start sites and alternative polyadenylation. As more differential splicing and sQTL detection methods are published, comparing the results of these various methods may provide a more nuanced assessment of the role of sQTLs in complex disease. Another limitation is that the GTEx v6 liver RNA-seq data was processed differently than the iPSC and HLC data, which may have resulted in the inability to detect the same sQTLs across cell types at the intron level. The use of primary liver tissue samples to interpret and replicate specific sQTLs we have cataloged here would be an important direction for future work.

In summary, sQTL discovery in iPSC-derived cells may serve as an approach complementary to primary tissue QTL discovery efforts and provide a model system in which to identify and functionally characterize sQTLs relevant for complex trait variation.

## Methods

### iPSC derivation and differentiation into HLCs

As previously described,^15^ iPSC cell lines were generated from the peripheral blood mononuclear cells (PBMCs) of 91 human subjects in good health and without cardiovascular disease. In brief, PBMCs were obtained from the subjects’ blood, cultured with cytokines, and expanded into erythroblasts. Erythroblasts were then transduced with Sendai viral vectors expressing Oct4, Sox2, Klf-4, and c-Myc. Once transduced, cells were transferred to culture containing mouse embryonic feeder cells. Colonies of iPSC cells were then moved to hESC medium until at least cell passage 12. Once established on the hESC medium, iPSC lines were removed from feeder culture and passaged at least 5 times before use in HLC differentiation. iPSC quality control procedures included mycoplasm testing, quantification of pluripotency markers,^25^ and confirmation of Sendai vector loss through RNA-sequencing. HLC differentiation from the iPSCs was conducted using feeder-free differentiation.^13^ RNA sequencing was used to confirm the expression of hepatocyte-specific genes in the HLC lines. Albumin, ApoB, and triglycerides were also assayed to confirm HLC quality.

### Genotyping and quality control

DNA was extracted from each donor with the QIAsymphony SP system (QIAGEN) and genotyped on the Infinium Human CoreExome-24 BeadChip (Illumina). Variants missing >5% of total genotypes and variants that deviated from Hardy-Weinberg equilibrium were removed. Haplotypes were phased using SHAPEIT2 and missing genotypes were imputed to the 1000 genomes phase 3 multi-ethnic panel. Following imputation, the same missingness and Hardy-Weinberg filters were applied to the imputed data.

### RNA sequencing data generation

As previously described,^15^ RNA was extracted from iPSC and HLC cells using the RNAeasy mini kit (QIAGEN) and assayed with the Agilent RNA 6000 Kit and Bioanalyzer. Libraries were prepared using TruSeq Stranded mRNA Library Prep Kit (Illumina). Sequencing was performed at the University of Pennsylvania’s Next Generation Sequencing center using Illumina Hi-Seq 2000/2500 Systems with 100bp/125bp paired-end reads. Target read depth for each sample was 50 million reads. We assessed quality of the raw FastQ files using FastQC^26^ and used TrimGalore!^27^ to trim adaptors. We then aligned trimmed reads that passed FastQC quality control to hg19/GRCh37 reference using the 2-pass mode of STAR aligner.^28^ Whole transcriptome RNA-sequencing data was generated for 89 iPSC and 86 HLC samples - we used the data from 83 individuals who had data from both iPSC and HLC lines and that passed differentiation and quality control standards.

### Splicing quantification

Alternative splicing events were quantified in both iPSC and HLC samples using Leafcutter 0.2.7 (http://davidaknowles.github.io/leafcutter/index.html, Accessed December, 2017), which uses split-mapped RNA-sequencing reads that span two different exons to identify excised introns. It then identifies groups of excised introns that share donor/acceptor splice sites. The proportion of reads that map to the individual introns within these intron groups (or “clusters”) are then used as quantitative estimates of alternative splicing. The intron clusters used in this analysis were identified using the Leafcutter: “python leafcutter_cluster.py -j juncfiles.txt -m 50 -o clusters -l 500000.”

### Differential splicing and expression analysis

We identified introns that are differentially spliced between iPSC and HLC samples using the differential splicing function of Leafcutter. The differential splicing analysis was then run using: “leafcutter_ds.R --num_threads 4 -- min_samples_per_group 10 --min_coverage 6 --exon_file=gencode19_exons.txt.gz clusters_perind_numers_noXY.counts.gz SVAconfounderMatrix.txt.” The gencode19_exons.txt.gz file is provided on the Leafcutter GitHub, and the SVA confounder matrix was generated using the SVA R package.^29^

Of the genes that were identified as differentially spliced, we conducted overrepresentation analysis of KEGG Pathway annotations using the R package ClusterProfiler.^30^ The background set of genes was comprised of all of the genes that were used for differential splicing analysis (i.e., was “successfully” tested for differential splicing analysis, in that it passed filtering thresholds). To compare iPSC and HLC samples based on patterns of alternative splicing, we conducted principal components analysis on the iPSC/HLC splicing phenotypes (normalized, standardized intron ratios) using the R package prcomp.

Using the DESeq2 R package^31^, we performed differential expression analysis between the 83 donor-paired iPSC and 83 HLC RNA-Seq samples on all transcripts that had at least 20 samples across all 166 RNA-Seq samples with 6 or more RNA-seq reads mapping. DESeq2 was run with a confounders matrix made up of 24 surrogate variables generated from the quantile normalized, standardized TPMs of the transcripts that met the previously mentioned expression thresholds. The surrogate variables were generated using the SVA R package.^29^ Transcripts were considered differentially expressed in the iPSCs or HLCs if they had a Benjamini and Hochberg adjusted p-value of ≤0.05 and a log_2_-fold-change of 2 or greater.

To compare iPSC and HLC samples based on patterns of gene expression, we also conducted principal components analysis on the iPSC/HLC normalized, standardized TPMs using the R package prcomp.

Finally, we used the ClusterProfiler^30^ R package to perform KEGG pathway enrichment analysis on the differentially expressed transcripts in both the HLC samples and the iPSC samples. The pathway enrichment analysis was done with a q-value cutoff of 0.05 and using a background set containing all transcripts that were tested for differential expression.

### sQTL identification

Clusters were identified within iPSC and HLC samples using the Leafcutter call: “python leafcutter_cluster.py -j juncfiles.txt -m 50 -o clusters -l 500000.” Intron proportions were standardized across individuals for each intron, and read ratios were quantile-normalized across introns using: “python prepare_phenotype_table.py perind.counts.gz -p1.” We tested for association between variants with MAF>0.05 and within 100kb of a given intron cluster using linear regression as implemented with QTLtools (v2-184, http://fastqtl.sourceforge.net/).^32^ To obtain the FDR <5% sQTLs used for the majority of the analyses, we used the “-permute 1000 10000 --grp-best” adaptive permutation command to obtain the most significant variant-intron pair per gene. sQTL scans for both iPSC and HLC samples were also run without permutations for use in qvalue, METASOFT, and colocalization analyses, since permuted results only output p-values for the most highly associated SNP per intron cluster/gene. In each scan we controlled for PEER factors, sex, and four genotype based principle components (calculated on the VCF of all genotypes filtered for variants MAF > 0.05 using EIGENSTRAT’s smartpca – genotypes were first LD pruned with PLINK using R^2^=0.8).^33,34^ PEER factors were estimated on the standardized, quantile-normalized splicing phenotypes matrix and were included as covariates in the regression. sQTL scans were run for both iPSC and HLC samples over a range of 1-20 PEER factors to determine the number of factors that maximized the numbers of sQTL discovered for each cell type. Five PEER factors were used in the HLC sQTL scan, while six PEER factors were used in the iPSC sQTL analysis. QTLtool’s adaptive permutations scan outputs p-values adjusted for the number of introns tested per gene. We then further corrected these p-values across genes using the Benjamini and Hochberg method to obtain a global false discovery rate.^35^

For the sQTL scan conducted in the GTEx v6 liver samples (n=96), a similar procedure was followed in order to obtain the nominal sQTLs used for the replication analysis. We used regression as implemented by FastQTL^36^ to obtain the nominal p-values used for replication qvalue analyses. Due to differences in data processing and ethnicity of the samples, 3 genotype PCs and 5 PEER factors were included as covariates.

### Enrichment analysis

#### Annotations

Splice site windows were obtained by taking Gencode v19 exon annotations and setting intervals 3bp into an exon and 8bp beyond an exon boundary to encompass the range of base pairs that encompass canonical binding sites for splicing machinery. Genomic regions were ascertained from Gencode v19 annotations using the GenomicFeatures R package.^37^ We retained known protein coding transcripts, and then obtained annotations for introns, 5’ UTR, 3’ UTR, coding sequence, and promoters using the intronsByTranscript, fiveUTRsByTranscript, threeUTRsByTranscript, cdsBy, and promoters functions, respectively. RNA binding protein peaks were downloaded from eCLIP files from ENCODE.^38^ Using the data from the same eCLIP paper that provided the ENCODE RBP files, we identified 33 RBPs implicated in either splicing regulation of 3’ processing. For each RBP, we took the intersection of each of the replicate peak files, then combined all of these intersected files to get one BED file of “RBP peaks.”

#### Enrichment analysis using GoShifter

GoShifter^39^ was used to determine the degree to which iPSC and HLC sQTLs (FDR <5%) were enriched in the annotations described above. We used LD files calculated with PLINK^34^ on our FDR <5% sQTLs to get the groups of SNPs in LD of R^2^>0.8 required for running GoShifter, which creates “circularized” loci and shifts annotations within these regions to provide a null distribution with which to compare the observed results. Odds ratios were calculated using the fisher.test function in R.

### Distance to 5’ and 3’ ends of genes and location within introns

sQTL variants were identified as being upstream, downstream, or within the excised intron with which they were associated. We then plot the distance of the variant to the 5’ or 3’ end of the intron cluster. If the sQTL variant fell in between the exons within the excised intron, intron length was divided by 10 and the sQTL was assigned to one of these deciles.

### Replication analysis in primary liver tissue

We used the R package qvalue^17,40^ to calculate the π_0_ values for the distribution of iPSC and HLC sQTL SNP-gene pairs in GTEx v6 liver data. For each of the SNP-gene pairs in the FDR < 5% iPSC and HLC sQTL scan, we pulled the nominal p-values for these same pairs out of the GTEx v6 liver sQTL scan to obtain an estimate of the replication rate.

### Cell-type specificity analyses

To evaluate the degree to which sQTLs are shared between cell-types, we used METASOFT^41^ to calculate the posterior probabilities (m-values) that an sQTL discovered in iPSC samples is also an sQTL in HLC samples, and vice-versa. An sQTL was considered associated with both cell types if the m-value was ≥0.9 for both cell-types. An sQTL was considered HLC-specific if the m-value was ≥0.9 in HLCs and <0.1 in iPSCs. Similarly, sQTLs were considered iPSC-specific if the m-value was ≥0.9 in iPSCs and <0.1 in HLCs.^42^ To facilitate comparisons between sQTLs and eQTLs, we conducted METASOFT analysis on the top one thousand sQTLs from our iPSC and HLC permutation scans. The same procedure was used for our iPSC and HLC eQTLs.

### eQTL identification

Aligned RNA-seq reads were assigned to Gencode v19 genomic annotations using the R package featureCounts.^43^ Transcripts were filtered for HLC and iPSC samples separately and were kept for analysis if at least 10 samples had 6 or more mapped RNA-seq reads. Raw featureCounts transcript counts were converted to TPMs. For each sample, the distribution of TPMs were normalized to the empirical average quantiles across samples using the R function normalizeQuantiles, and then, for each gene, for each cell type, the distribution of TPMs was transformed to the quantiles of the standard normal distribution, using the R function qqnorm. We tested for association between bi-allelic variants with MAF>0.05 and within 1Mb of a given transcript (beginning or end of transcript) using linear regression as implemented with QTLtools (v2-184, http://fastqtl.sourceforge.net/). To obtain the FDR <5% eQTLs used for the majority of the analyses, we used the “--permute 1000 10000” adaptive permutation command to get the most significant variant-gene pair. eQTL scans for both iPSC and HLC samples were also run without permutations for use in METASOFT and colocalization analyses, since permuted results only output p-values for the most highly associated SNP per gene.

As with the sQTL analysis, in each eQTL scan we controlled for PEER factors^44^, gender, and four genotype based principle components (calculated using EIGENSTRAT’s smartpca, and first LD pruned R^2^=0.8 with PLINK).^33,34^ PEER factors were estimated using on the standardized, quantile-normalized TPMs calculated from the featureCounts output and were included as covariates in the regression. eQTL scans were run for both iPSC and HLC samples over a range of 1-30 PEER factors to determine the number of factors that maximized the numbers of eQTLs discovered for each cell type. 20 PEER factors were used in the HLC eQTL scan, while 16 PEER factors were used in the iPSC eQTL scan. QTLtools’ adaptive permutation scan outputs p-values adjusted for the number of variants tested per gene. We then further corrected these p-values across genes using the Benjamini and Hochberg method to obtain a global FDR.^35^

### Credible Set Calculations

To calculate 95% credible sets for each of the iPSC/HLC FDR < 5% sQTLs, we used the process described in Maller et al. 2012 to identify sets of variants with a 95% probability of containing the causal variant.^45^

### Colocalization analyses

Using the default parameters for the R package coloc^20^ (CRAN v3.1) we conducted colocalization analysis between each of the GWAS traits from the Global Lipids Genetics Consortium^46^ (HDL-C, LDL-C, TG, and TC) and the FDR <5% eQTL-gene and sQTL-gene pairs from our HLC samples. For each GLGC trait, we ranked genome-wide significant variants by p-value and chose the most significant variants within 100kb windows of each other. If any of the FDR <5% eQTL/sQTL variants fell within 100kb of these GLGC variants, we conducted colocalization analysis on all variants within 100kb of the GWAS variants, using the p-values obtained from our nominal sQTL/eQTL scans. We used the MAF estimated from the GLGC.

---

**Supplementary Figure 1.**
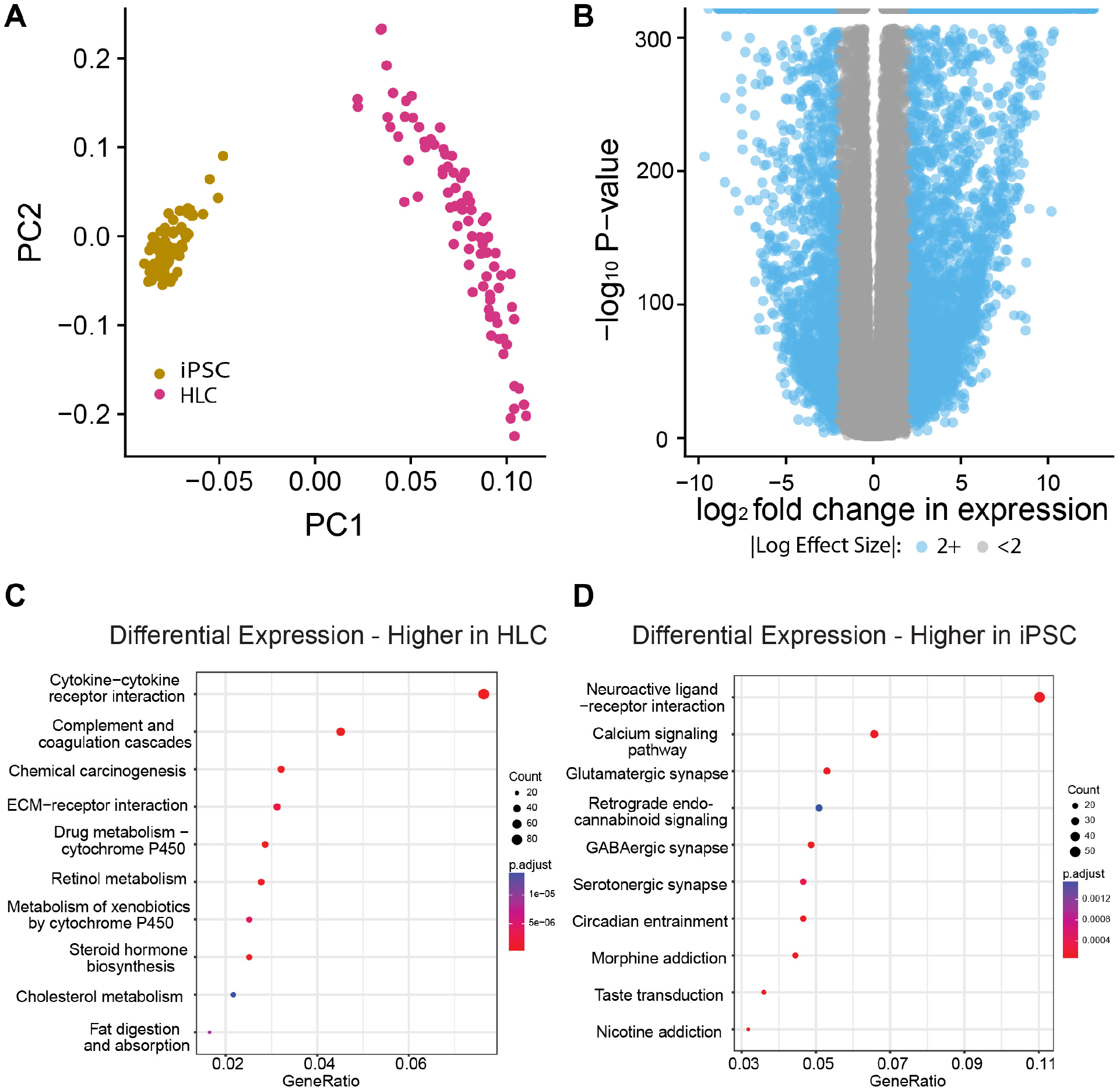
Genes that are differentially expressed between iPSC and HLC samples (FDR <5%): a) PCA of normalized TPMs, b) Volcano plot of differentially spliced genes (FDR <5%), c) KEGG pathway enrichment analysis for genes that are differentially expressed more highly in HLC relative to iPSC, d) KEGG pathway enrichment analysis for genes that are differentially expressed less highly in HLC relative to iPSC.

**Supplementary Figure 2.**
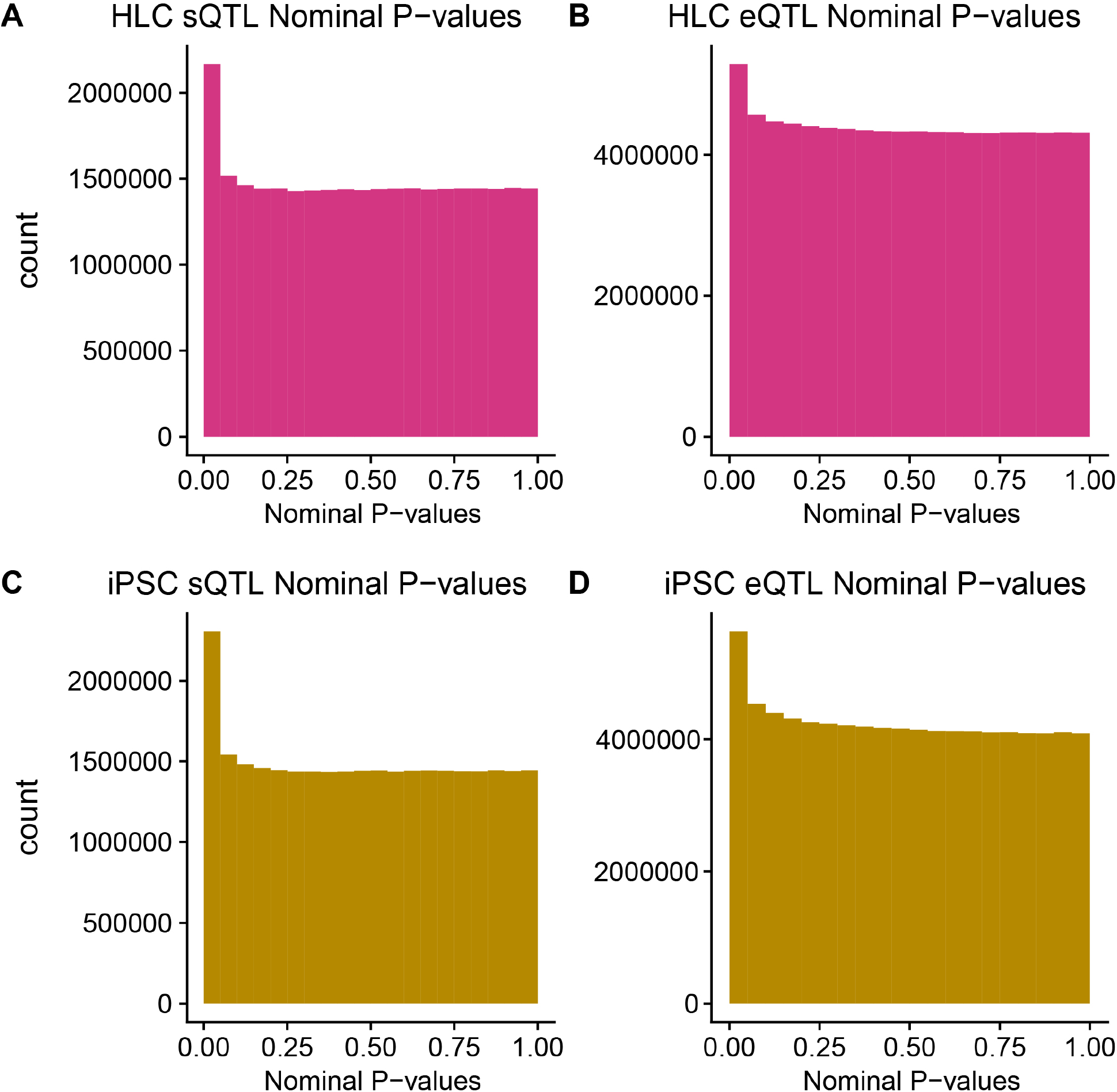
P-value histograms for nominal sQTL and eQTL scans.

**Supplementary Figure 3.**
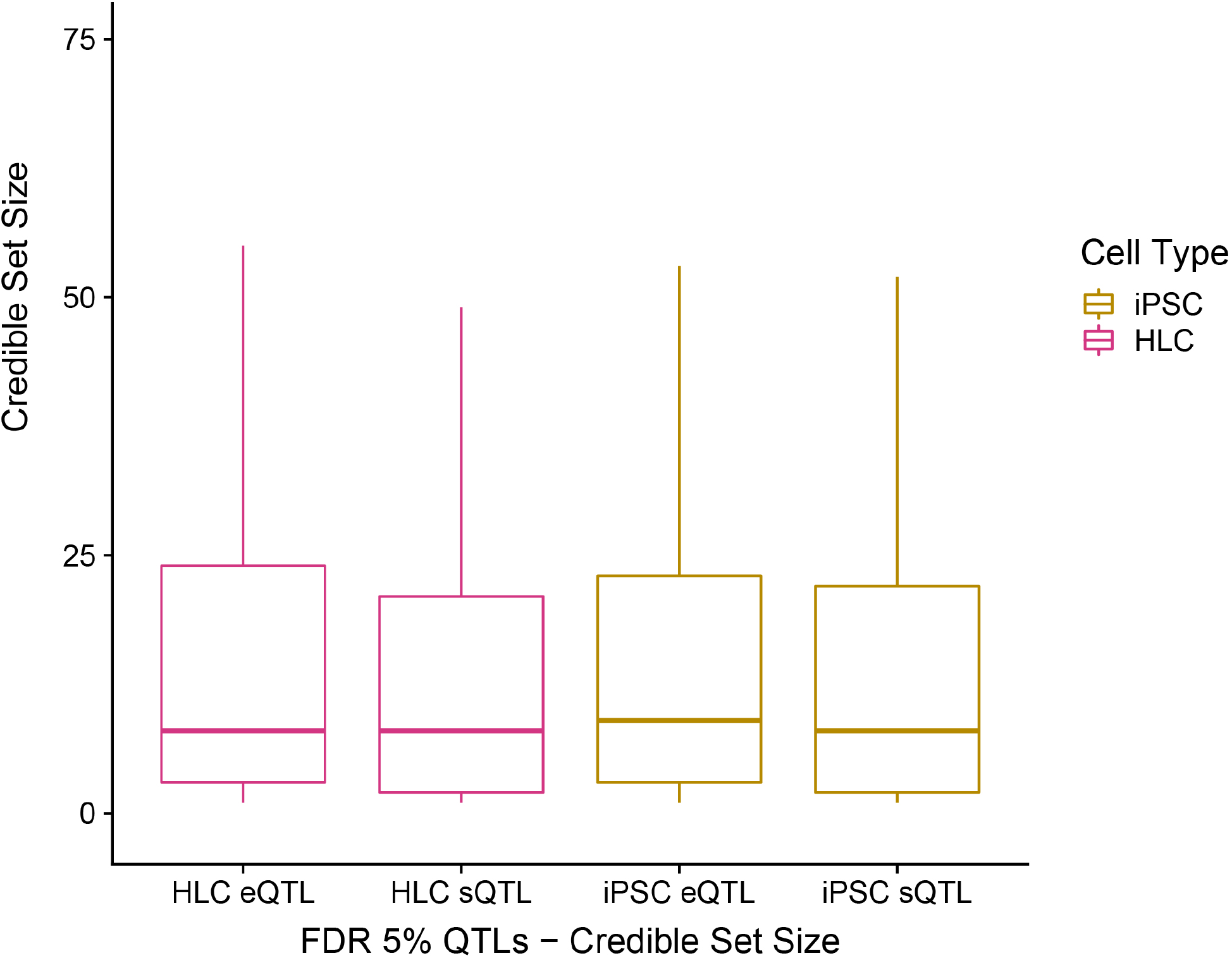
Credible sets for iPSC, HLC sQTLs and eQTLs. Boxplot whiskers calculated as 1.5 * interquartile range. Not pictured: 62 sets for iPSC sQTLs, 200 sets for iPSC eQTLs, 73 sets for HLC sQTLs, 116 sets for HLC eQTLs.

